# Effects of memantine and high dose vitamin D on gait in the APP/PS1 mouse model of Alzheimer’s disease following vitamin D deprivation

**DOI:** 10.1101/2021.09.21.461069

**Authors:** Dana Broberg, Dickson Wong, Miranda Bellyou, Manuel Montero-Odasso, Olivier Beauchet, Cedric Annweiler, Robert Bartha

## Abstract

**Background:** Altered gait is a frequent feature of Alzheimer’s disease (AD), as is vitamin D deficiency. Treatment with memantine and vitamin D can protect cortical axons from exposure to amyloid-β and glutamate toxicity, suggesting this combination may mitigate altered gait in AD.

**Objective:** Investigate the effects of vitamin D deprivation and subsequent treatment with memantine and vitamin D enrichment on gait performance in APPswe/PS1dE9 mice.

**Methods:** Male APPswe/PS1dE9 mice were split into four groups (*n*=14 each) at 2.5 months of age. A control group was fed a standard diet throughout while the other three groups started a vitamin D-deficient diet at month 6. The VitD− group remained on this deficient diet for the rest of the study. At month 9, the remaining two groups began treatment with either memantine alone or memantine combined with 10 IU/g of vitamin D. Gait performance was assessed at months 6, 9, 12, and 15.

**Results:** Vitamin D deprivation led to a 13% increase in hind stride width by month 15 (*p*<0.001). Examination of the treatment groups at month 15 revealed that mice treated with memantine alone still showed an increase in hind stride width compared to controls (*p*<0.01), while mice treated with memantine and vitamin D did not (*p*=0.21).

**Conclusion:** Vitamin D deprivation led to impaired postural control in the APPswe/PS1dE9 model. Treatment with memantine and vitamin D, but not memantine alone, prevented this impairment. Future work should explore the potential for treatments incorporating vitamin D supplementation to improve gait in people with AD.

## INTRODUCTION

Alzheimer’s disease (AD) is a neurodegenerative disease characterized by the development of amyloid-β plaques and neurofibrillary tangles coupled with progressive cognitive decline. AD has widespread prevalence in the elderly, affecting one-third of individuals over the age of 85 [1]. Motor abnormalities (e.g. gait disorders) frequently occur in AD patients [2], often prior to the onset of cognitive decline [3]. Gait disorders cause reduced mobility, thereby decreasing functional independence and quality of life [4–6]. More specifically, individuals with AD tend to walk with reduced pace, shorter stride length and, in some cases, larger stride width [3].

Potentially compounding this problem, it is estimated that as many as 70-90% of individuals with AD have deficient levels of vitamin D [7]. Many studies suggest that vitamin D has neuroprotective effects in AD, which include neurotransmitter regulation [7, 8], prevention of calcium toxicity [7, 8], immune system regulation [7–9], induction of amyloid-β clearance [7–10], prevention of oxidative stress [8], and neurotrophic factor regulation [7–10]. As such, vitamin D deficiency is associated with increased risk of cognitive decline in individuals with AD [7, 9, 11, 12], which may also contribute to worsening motor ability [13].

Recently, the use of memantine, an *N*-methyl-*D*-aspartic acid (NMDA) receptor antagonist approved for the symptomatic treatment of moderate to severe AD, in combination with vitamin D supplementation has been investigated as a treatment for AD. This treatment combination can protect cortical axons from exposure to amyloid-β and glutamate toxicity [14]. The combined effects of memantine and vitamin D have also shown greater potential to slow cognitive decline in AD patients compared to either memantine or vitamin D alone [15]. Furthermore, both memantine and vitamin D have been shown to improve physical performance and gait in AD patients when taken separately [16–18]. Therefore, the combination of memantine and vitamin D may elicit greater improvements in gait compared to treatment with either alone.

Transgenic mouse models of AD are frequently used to study the mechanisms underlying disease progression as well as treatment efficacy. The severity of AD pathology varies across these models based on the degree of genetic manipulation, such that mice containing a higher number of AD-related genetic mutations result in more aggressive, or fast-progressing models. The double transgenic APPswe/PS1dE9 model is considered a preclinical or mild model of AD, reflecting changes in the early stages of the disease. Specifically, APPswe/PS1dE9 mice present with amyloid plaques starting after six months [19], neuronal loss at nine months [20], and evidence of cognitive impairment by 12 months [21]. Neurofibrillary tangles are absent in this model. These mice also display progressive loss of motor coordination in wire hanging, rotarod and balance beam tests [22, 23]. However, spatiotemporal parameters of gait have yet to be characterized in this model. Due to the slow onset of amyloid pathology, the APPswe/PS1dE9 model is well-suited to study changes in gait over time.

The specific objectives of this study were to assess the effects of vitamin D deprivation on gait performance in the APPswe/PS1dE9 mouse model of AD and then determine whether treatment with memantine, or memantine combined with vitamin D enrichment reversed changes in gait. Gait performance was measured by the CatWalk Gait Analysis System (Noldus Information Technology, Wageningen, The Netherlands). We hypothesized that vitamin D deficiency would lead to altered spatiotemporal parameters of gait, while the combination of memantine plus vitamin D supplementation in vitamin D deficient mice would reverse gait decline.

## MATERIALS AND METHODS

### Animal Subjects

A description of the mice used in this study has been previously published [24]. Briefly, 60 male APPswe/PS1dE9 mice were obtained from Jackson Laboratories (JAX MMRC Stock #034829; Bar Harbour, ME, USA) at 2.5 months of age. These mice carried the Swedish mutation of the human amyloid precursor protein (APPswe) and the exon-9 deleted mutation of the human presenilin-1 protein (PS1dE9) on a (C57BL/6 x C3H) F2 genetic background. Mice arrived in cohorts of 5 until a total of 60 mice were obtained. The mice were housed individually in standard tub cages with *ad libitum* access to food and water, and a covered shelter as the only form of environmental enrichment. A 12:12 h light-dark cycle was maintained, with lights on at 7 am. All experimental procedures were completed between the hours of 9 am and 6 pm. Mice were euthanized at 15 months of age. Ethics approval for this study was obtained from the University of Western Ontario Animal Use Subcommittee (AUP#: 2012-040).

### Diet and Serum Vitamin D Levels

A summary of the study design is shown in Figure 1. The scheme used to modify vitamin D levels was published previously [24] but is included here for completeness. From 2.5 to 6 months of age, all 60 mice were fed a standard AIN-76A rodent diet from Research Diets, Inc. (D10001; New Brunswick, NJ, USA). This control diet contained 1000 IU vitamin D_3_ per kg. At 6 months of age, the surviving 56 mice were randomized to one of four groups. The control group continued to receive the control diet for the remainder of the study. There were 14 mice in the control group at 6 and 9 months of age, and 13 mice at 12 and 15 months of age (Control, *n*=14, 14, 13, 13). The other three groups (*n*=42) received a vitamin D-deficient diet from Research Diets, Inc. (D08090903; New Brunswick, NJ, USA) with <1 IU vitamin D_3_ per kg from 6 to 9 months of age. Other than the omission of vitamin D from the diet’s vitamin mix, the vitamin D-deficient diet was identical to the standard AIN-76A diet. The vitamin D-deficient group (VitD−, *n*=14, 14, 13, 13) continued to receive the vitamin D-deficient diet for the remainder of the study. Starting at month 9, the remaining two groups were fed 20-30 mg/kg per day of memantine in addition to either the vitamin D-deficient diet (Mem & VitD−, *n*=14, 13, 13, 13), or a vitamin D-enriched diet containing 10,000 IU vitamin D_3_ per kg of food (Mem & VitD+, *n*=14, 14, 14, 14). These diets were also provided by Research Diets, Inc. (D16030202 and D16030203, respectively; New Brunswick, NJ, USA). As described previously [24], this high vitamin D dose would have been comparable to a dose of 6700 IU/day in humans. This high dose was not only postulated to be necessary for neuroprotection [25] but was also proven to be safe in a recent clinical trial [26]. The memantine dose was chosen based on previous studies showing that treatments using 20-30 mg/kg per day resulted in plasma drug levels of ∼1 μM [27, 28], which both preclinical and clinical studies indicate is therapeutic [29, 30].

**Figure 1.**
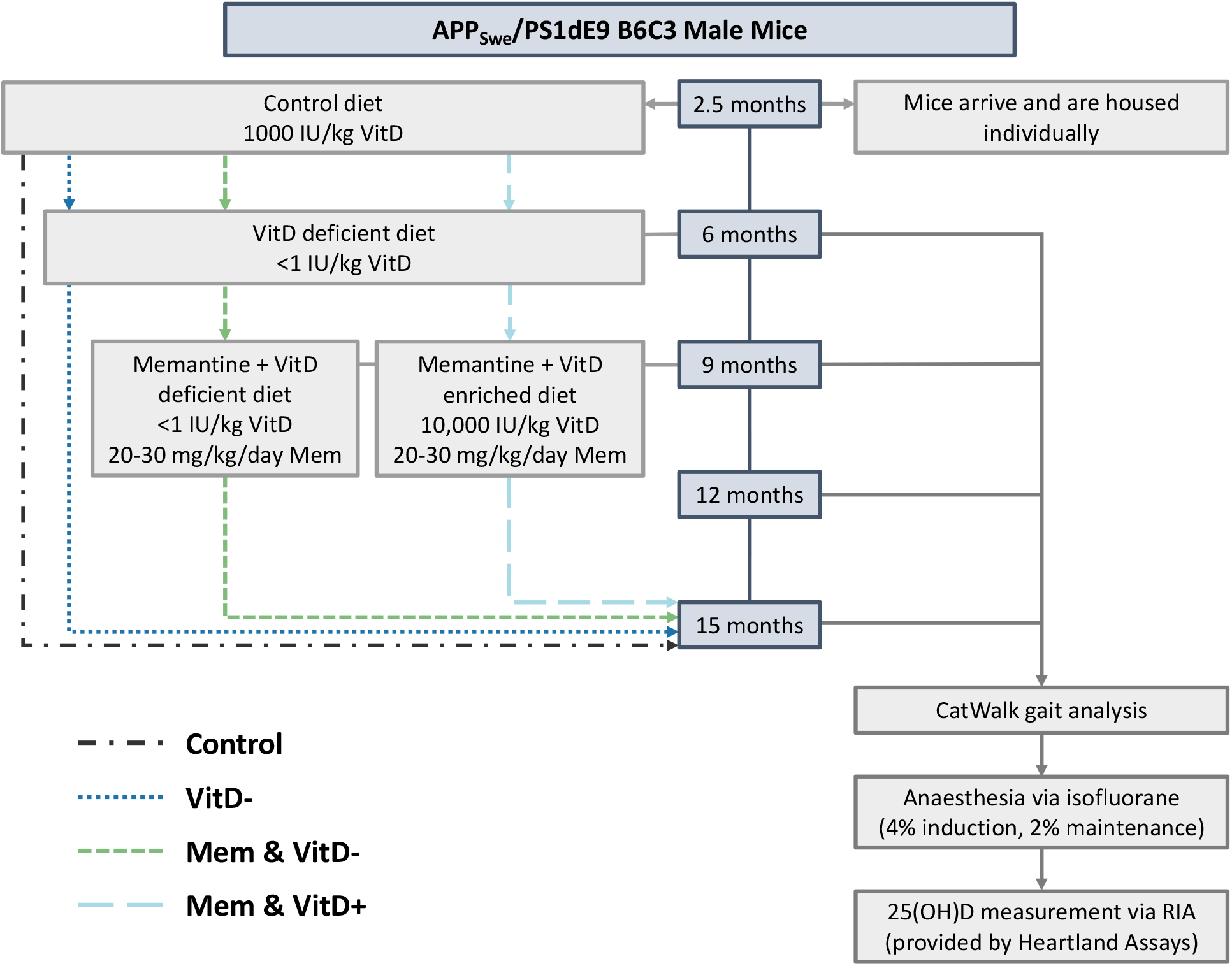
Summary of the study design: Changes in diet at months 6 and 9 were implemented *after* measurements of gait and serum 25(OH)D. VitD−, vitamin D-deficient; Mem, memantine; VitD+, vitamin D-enriched.

To ensure that the vitamin D levels in the mice were modified by the diet, mice were anaesthetized using isofluorane gas (4% induction, 2% maintenance) and a tail nick procedure was performed for blood draw at months 6, 9, 12, and 15. This procedure was performed after the CatWalk tests to ensure that the anaesthesia did not affect the gait analysis. Blood samples were then sent to Heartland Assays (Ames, IA, USA) for measurement of serum 25(OH)D levels by radioimmunoassay.

### CatWalk Gait Analysis

Gait analysis was performed at 6, 9, 12, and 15 months of age (Fig 1) using CatWalk Automated Gait Analysis System version 7.1 (Noldus Information Technology, Wageningen, The Netherlands), which has been described in detail elsewhere [31]. Briefly, mice were individually placed into the walkway and allowed to freely cross the enclosed glass plate which was illuminated by fluorescent light. Contact of the mouse with the glass plate reflected light downwards to a high-speed colour camera, allowing the detection of each footprint. The camera was connected to a computer equipped with the CatWalk acquisition software. Runs where the mouse crossed the walkway at a constant, moderate speed were considered successful. For each mouse and time point, multiple runs were performed until at least two successful runs were recorded. Analysis was then performed on the best two runs. Footprints were manually labelled as right front, right hind, left front, or left hind. The CatWalk software was used to quantify several spatiotemporal parameters of gait including swing, stride length, swing speed, and stride width, which are described in Table 1. The CatWalk software provided the mean and standard deviation for the swing, stride length, and swing speed of each limb separately. Therefore, for each of these gait parameters, a pooled mean and standard deviation from all four limbs was calculated for each run. As a measure of regularity, the coefficient of variation (CoV; standard deviation/mean) was calculated for swing, stride length and swing speed. CatWalk did not provide standard deviation for front or hind stride width and therefore CoV could not be calculated for these two gait measures. The swing, CoV of swing, stride length, CoV of stride length, swing speed, CoV of swing speed, front stride width, and hind stride width of the two best runs for each mouse and time point were averaged and statistically compared between groups.

**Table 1.** CatWalk gait parameter descriptions

| *Parameter* | *Description* |
| --- | --- |
| Swing (s) | Duration without contact with the glass plate in a step cycle |
| Stride Length (mm) | Distance between successive placements of the same paw |
| Swing Speed (m/s) | Speed of the paw during a swing |
| Stride Width (mm) | Average width between either the front paws or the hind paws (referred to as Base of Support by CatWalk) |

### Statistical analysis

All statistical analyses were performed using GraphPad Prism Version 9.0 for MacOS (GraphPad Software, San Diego, CA). Outliers were identified using the ROUT method [32]. Mice identified as outliers in serum 25(OH)D levels at a specific time point were removed from 25(OH)D analyses at that time point. Mice determined to be outliers for gait measures at a specific time point were also removed from all gait analyses at that time point.

Both the serum vitamin D levels and individual gait measurements were analyzed using a mixed-effects model, as implemented in GraphPad Prism 9.0, which fits a compound symmetry covariance matrix using Restricted Maximum Likelihood (REML). In the presence of unbalanced data (unequal sample sizes), as was the case for these measurements, results can be interpreted like repeated measures two-way ANOVA (which is incapable of handling missing values). Age, treatment, and age-by-treatment interaction were considered fixed effects. Homoscedasticity and normality were confirmed using residual plots and QQ plots, respectively. Sphericity was not assumed, so the Geisser-Greenhouse correction was applied [33]. When any fixed effect was found to be significant, post-hoc tests were performed to identify statistically significant differences between time points and treatment groups. In all post-hoc tests, multiplicity adjusted *p*-values were computed. Significance for all statistical tests was set at *p* < 0.05.

## RESULTS

### Serum Vitamin D levels

The number of mice included in the 25(OH)D analysis in each treatment group at months 6, 9, 12 and 15, respectively, was as follows: control, *n*=14, 14, 13, 13; VitD−, *n*=14, 13, 10, 7; Mem & VitD−, *n*=14, 12, 13, 13; Mem & VitD+, *n*=14, 14, 14, 8. 18 mice at various time points were excluded because their samples were lost during transit to Heartland Assays. The sample loss was random; therefore, these mice were still included at other time points for which their serum levels were available. The comparison of 25(OH)D levels in these groups has been published previously [24] but are described here briefly for completeness. Mice on the control, vitamin D-deficient, and vitamin D-enriched diets had mean serum 25(OH)D levels of 27, 3, and 61 ng/ml, respectively. There were no significant changes in serum 25(OH)D in the control group throughout the study. In contrast, the switch from the control diet to the vitamin D deficient diet at month 6 resulted in an average 83% decline in 25(OH)D levels by month 9 that declined further at months 12 and 15. As expected, the switch from a vitamin D deficient diet to a vitamin D enriched diet at month 9 in the Mem & VitD+ group resulted in a large increase in serum 25(OH)D: at months 12 and 15, 25(OH)D levels in this group were more than doubled compared to month 6. Therefore, the enriched diet not only recovered the mice from vitamin D deficiency but increased their serum vitamin D levels above the level of the control diet. Overall, these results indicate that serum vitamin D levels of the mice were modified by the diets as intended. Mice on the vitamin D enriched diet had more than 10× higher serum 25(OH)D levels compared to the mice in the vitamin D deficient diet.

### Gait Analysis – Effect of Vitamin D Deprivation

The number of mice included in the gait analysis in each group at months 6, 9, 12 and 15, respectively, was as follows: control, *n*=14, 14, 13, 13; VitD−, *n*=12, 13, 13, 10; Mem & VitD−, *n*=14, 13, 13, 13; Mem & VitD+, *n*=13, 14, 10, 14. CatWalk was used to determine several spatiotemporal parameters of gait in each treatment group throughout the study. To investigate the effects of vitamin D deprivation on gait in AD, we compared the gait of the control and VitD− treatment groups at each time point of the study. Measurements made at month 6 act as a baseline, as both groups had been fed a control diet up to this time point.

Presented first are the gait parameters for which a coefficient of variation was available: the swing, stride length, swing speed (Fig 2). The linear mixed model for swing revealed a significant effect of age (*F*(2,54) = 3.74, *p* < 0.05), but not treatment (*F*(1,26) = 0.77, *p* = 0.39) or age-by-treatment interaction (*F*(3,68) = 0.61, *p* = 0.61). Post-hoc comparison tests showed that swing significantly increased from month 12 to month 15 in the control group (14% increase, *p* < 0.05). Within the VitD− group, swing tended to increase from month 9 to month 15, however this was not statistically significant (17% increase, *p* = 0.08). Additionally, there were no significant differences between other time points in either group. The linear mixed model for swing CoV revealed a significant effect of age-by-treatment interaction (*F*(3,68) = 2.89, *p* < 0.05), but not treatment (*F*(1,26) = 0.84, *p* = 0.37) or age (*F*(2,58) = 0.79, *p* = 0.49). Post-hoc tests showed that swing CoV was significantly lower in the VitD− group versus the control group at month 9 (16% decrease, *p* < 0.05), however no differences were seen at other time points. Overall, vitamin D deprivation did not have a clear effect on swing or swing regularity.

**Figure 2.**
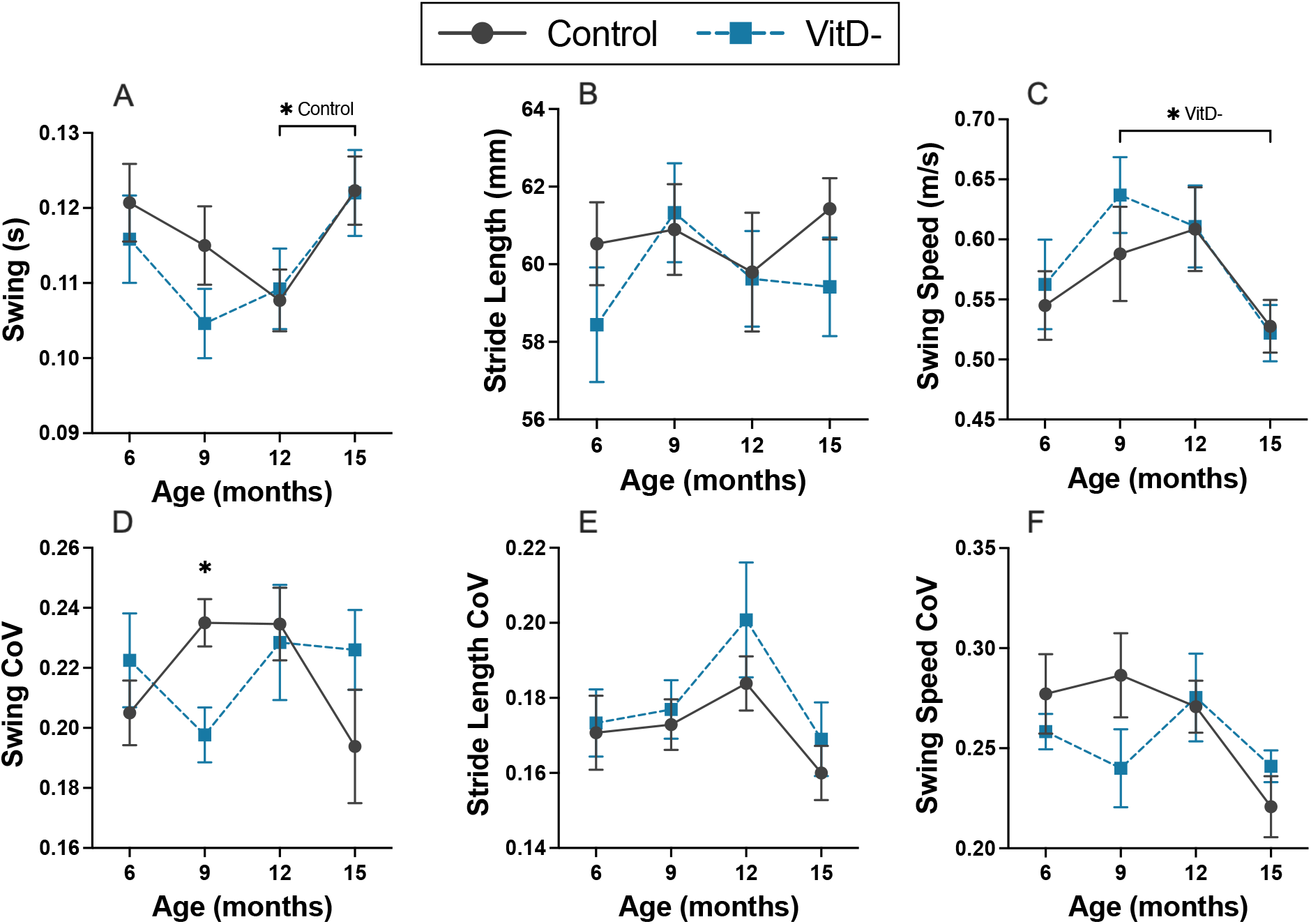
Effect of vitamin D deprivation on swing, stride length, and swing speed: The CatWalk gait analysis system was used to measure spatiotemporal parameters of gait in APPswe/PS1dE9 mice at 6, 9, 12 and 15 months of age. Mice in the ViD− group received a vitamin D-deficient diet after the six-month time point measurement. Shown here are swing (A), stride length (B), swing speed (C) and their respective coefficients of variation (D-F) in the control and VitD− groups. CoV; coefficient of variation; VitD−, vitamin D-deficient. \**p* < 0.05.

The linear mixed model for stride length revealed there were no significant effects of age (*F*(2,58) = 0.79, *p* = 0.49), treatment (*F*(1,26) = 0.86, *p* = 0.36), or age-by-treatment interaction (*F*(3,68) = 0.60, *p* = 0.62). The linear mixed model for stride length CoV revealed a significant effect of age (*F*(2,56) = 3.81, *p* < 0.05), but not treatment (*F*(1,26) = 0.84, *p* = 0.37) or age-by-treatment interaction (*F*(3,68) = 0.27, *p* = 0.85). Stride length CoV tended to increase from month 6 to month 12, but then decreased at month 15. Post-hoc comparisons, however, showed no significant changes in stride length CoV with age within treatment groups. In summary, vitamin D deprivation did not influence stride length or stride length regularity.

The linear mixed model for swing speed revealed a significant effect of age (*F*(2,55) = 3.54, *p* < 0.05), but not treatment (*F*(1,26) = 0.45, *p* = 0.51) or age-by-treatment interaction (*F*(3,68) = 0.29, *p* = 0.84). Post-hoc tests showed that the swing speed of the VitD− group was significantly decreased from month 9 to month 15 (18% decrease, *p* < 0.05). The linear mixed model for swing speed CoV revealed there were no significant effects of age (*F*(2,79) = 2.18, *p* = 0.11), treatment (*F*(1,94) = 0.66, *p* = 0.42), or age-by-treatment interaction (*F*(3,94) = 1.37, *p* = 0.26). Overall, vitamin D deprivation did not modify swing speed or swing speed regularity.

Next, front and hind stride widths were considered (Fig 3). CatWalk does not provide standard deviation for these measures, and therefore we could not calculate coefficient of variation for these two metrics. The linear mixed model for front stride width revealed there were no significant effects of age (*F*(2,66) = 0.37, *p* = 0.77), treatment (*F*(1,26) = 4.05, *p* = 0.054), or age-by-treatment interaction (*F*(3,68) = 2.56, *p* = 0.06). However, the linear mixed model for hind stride width revealed a significant effect of treatment (*F*(1,26) = 8.57, *p* < 0.01), but not age (*F*(2,55) = 0.48, *p* = 0.66) or age-by-treatment interaction (*F*(3,68) = 2.12, *p* = 0.11). Post-hoc comparison tests showed that the hind stride width of the VitD− group was significantly higher than that of the control group by month 15 (13% increase, *p* < 0.001). Therefore, vitamin D deprivation resulted in increased hind stride width but had no effect on front stride width.

**Figure 3.**
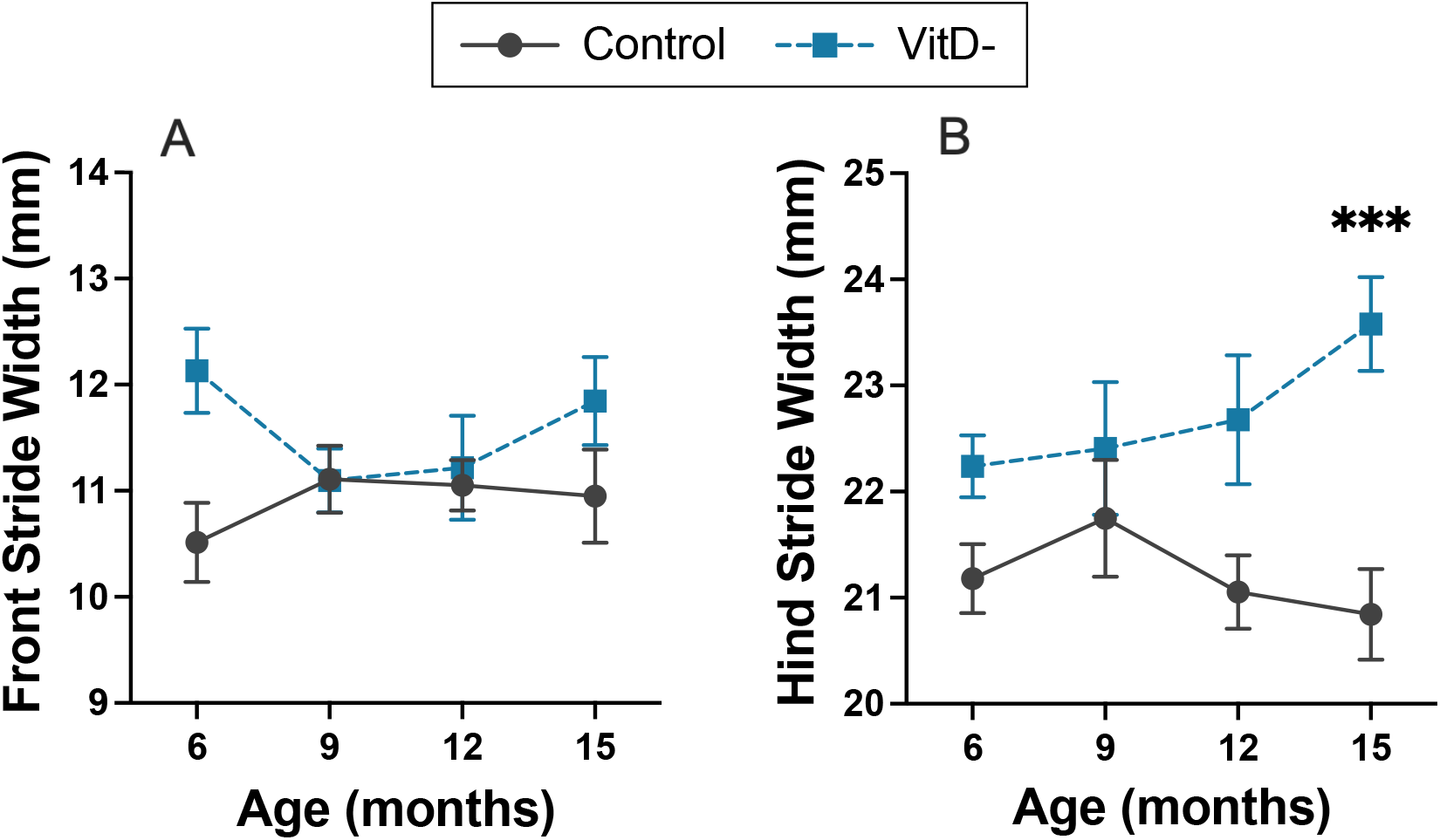
Effect of vitamin D deprivation on front and hind stride width: The CatWalk gait analysis system was used to measure spatiotemporal parameters of gait in APPswe/PS1dE9 at 6, 9, 12 and 15 months of age. Mice in the VitD− group received a vitamin D-deficient diet after the six-month time point measurement. Shown here are front (A) and hind (B) stride width in the control and VitD− treatment groups. VitD−, vitamin D-deficient. \*\*\**p* < 0.001.

### Gait Analysis – Effect of Memantine & Vitamin D on Vitamin D-Deficient Mice

Given that the only significant effect of vitamin D deprivation was an increase in hind stride width at month 15, the effect of memantine and vitamin D treatments on this parameter were evaluated at month 15. Specifically, a one-way ANOVA was performed that included all four groups to evaluate the effect of treatment on hind stride width at month 15 (Fig 4). The effect of treatment was statistically significant (*F*(3,47) = 8.06, *p* < 0.001). Animals treated with memantine alone that remained vitamin D deficient (Mem & VitD−) still had significantly higher hind stride width compared to controls (9% increase, *p* < 0.01). In addition, there was no difference in hind stride width between the VitD− group and the Mem & VitD− group (*p* = 0.61). These results suggest that memantine alone could not reverse the effects of vitamin D deficiency. In contrast, the Mem & VitD+ group had the same hind stride width as controls (*p* = 0.21) and, more importantly, lower hind stride width compared to the VitD− group (7% decrease, *p* < 0.05). Therefore, treatment with memantine and high dose vitamin D, but not memantine alone, was able to recover the vitamin D-deprived APPswe/PS1dE9 mice from the increase in hind stride width.

**Figure 4.**
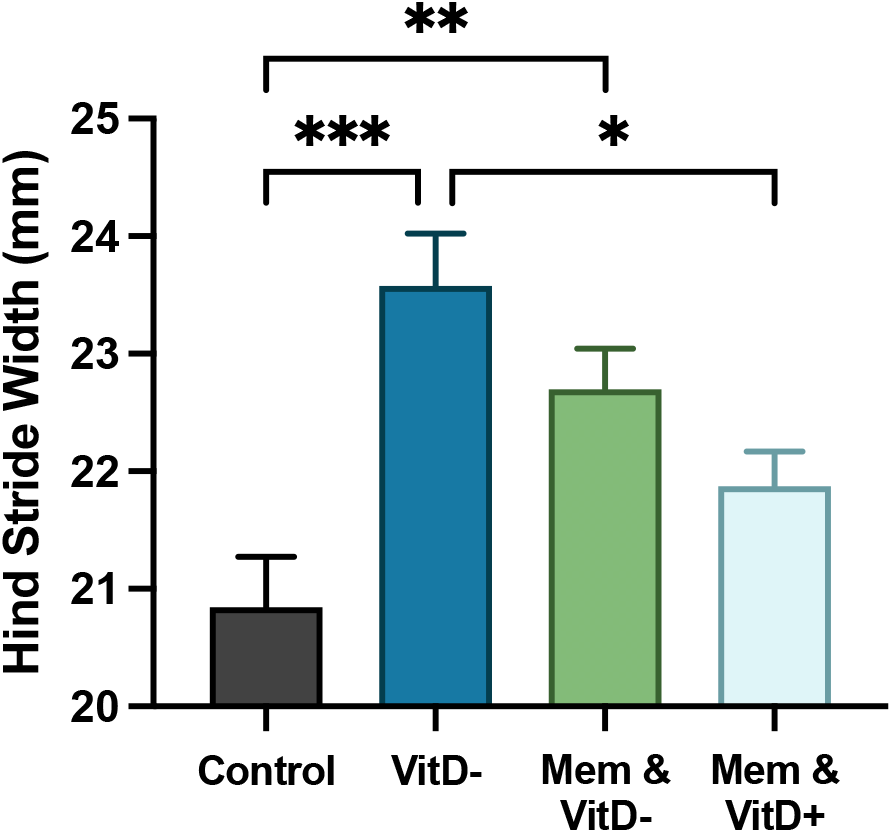
Effect of memantine plus vitamin D on hind stride width of vitamin D deficient mice: The CatWalk gait analysis system was used to measure hind stride width in APPswe/PS1dE9 mice. All groups were fed a vitamin D-deficient diet starting at six months. Treatment began for the Mem & VitD− and Mem & VitD+ groups at nine months and continued for the duration of the study. Shown here is hind stride width of the Mem & VitD−, Mem & VitD+, and VitD− treatment groups at 15 months of age. Mem, memantine; VitD+, vitamin D-enriched; VitD−, vitamin D-deficient. \**p* < 0.05, \*\**p* < 0.01, \*\*\**p* < 0.001.

For completeness, longitudinal analyses of gait of the VitD−, Mem & VitD−, and Mem & VitD+ treatment groups from month 9 onwards are provided in the Supplementary Materials. In these analyses, measurements made at month 9 were considered a baseline, as all three groups had been fed the same diets and were vitamin D deficient at this age. To summarize the results of this longitudinal analysis, age had a significant effect on swing, swing speed, and swing speed regularity; age-by-treatment interaction was significant for stride length regularity; and treatment alone did not have a significant effect on the gait parameters.

## DISCUSSION

This study examined the effects of vitamin D deprivation on gait in the APPswe/PS1dE9 mouse model of AD as well as subsequent treatment with memantine and vitamin D enrichment. The vitamin D deficient diets caused a decrease in serum vitamin D while the enriched diet caused an increase in serum vitamin D above the level of the control group, whose serum 25(OH)D level did not change throughout the study. Vitamin D deprivation led to increased hind stride width by month 15, but no changes in swing, swing regularity, stride length, stride length regularity, swing speed, swing speed regularity, or front stride width. Treatment with memantine and vitamin D enrichment, but not memantine alone, prevented the increase in hind stride width observed in vitamin D-deprived mice.

### Effect of Vitamin D Deprivation

The present study is the first to examine spatiotemporal gait parameters in the APPswe/PS1dE9 mouse model and the impact of vitamin D deprivation. Contrary to our hypothesis, we found that vitamin D deprivation did not modify swing, swing regularity, stride length, stride length regularity, swing speed, swing speed regularity, or front stride width. These results differ from a previous study in C57BL/6 J mice which found that mice fed a vitamin D-insufficient diet starting at 1 month of age exhibited significantly shorter stride length by 8 and 12 months of age [34]. However, the authors attributed this finding to reduced muscle mass, which we have previously shown was not observed in the mice used in the current study [24]. The current study also differed in the time of initiation and the duration of vitamin D deprivation. It is also possible that disturbances of balance control precede disturbances in displacement [35].

Interestingly, the current study showed that vitamin D deficiency led to an increase in hind stride width, a marker of balance and postural control [3], by month 15 of the study. We have previously shown that vitamin D deficiency does not cause any changes in body composition in APPswe/PS1dE9 mice [24], however it is possible that vitamin D deficiency affected gait through neurological processes. Another study in APPswe/PS1dE9 mice found progressive development of morphological abnormalities in the cerebellar cortex resulting in decreased performance in wire-hanging and rotarod tests [22]. As vitamin D deficiency is known to increase the risk for AD-associated cognitive decline [7], it is possible that vitamin D deficiency in the present study accelerated the development of cerebellar abnormalities, leading to the observed increase in hind stride width compared to mice fed a control diet. The current study suggests that vitamin D deficiency may lead to worsened gait in Alzheimer’s disease. Though further investigation is required in humans, already, we can see the potential value of maintaining serum 25(OH)D levels within the normal range. In humans, serum 25(OH)D levels greater than 20 ng/ml are often considered adequate [36]. For individuals over the age of 70, the recommended daily intake of vitamin D to achieve adequate serum 25(OH)D levels is 800 IU/day [36].

### Effect of Memantine & Vitamin D on Vitamin D-Deficient Mice

Following the observed increase in hind stride width due to vitamin D deficiency, we investigated the effects of treatment with memantine combined with vitamin D enrichment and memantine alone on hind stride width in mice that were vitamin D deficient. Interestingly, the combination treatment, but not memantine alone, was able to prevent increased hind stride width in vitamin D-deficient mice. Therefore, the combination of memantine and high dose vitamin D was an effective treatment for this gait abnormality in our vitamin D deficient APPswe/PS1dE9 model. This result is consistent with human studies of memantine and vitamin D supplementation in AD, which have both been found to improve measures of physical performance [16–18]. Memantine, in particular, is thought to regulate gait at a higher level of control, as it has been shown to reduce stride time variability in individuals with AD without affecting mean stride time [18].

It seems sensible that vitamin D enrichment was necessary for a reversal in the increased hind stride width caused by vitamin D deprivation. However, it is imperative to distinguish between normal vitamin D intake and high dose vitamin D enrichment. The current treatment did not simply restore serum vitamin D levels to that of control mice, but rather exceeded those levels by a factor greater than two. As described previously [24], the mice in this study received a vitamin D dose that would be comparable to humans receiving 6,700 IU/day. Research suggests that such a high dose may be necessary to achieve neuroprotective effects through antioxidative mechanisms, neuronal calcium regulation, and immunomodulation [25]. We hypothesize that it is through these mechanisms and their effects on motor cognition that gait was improved in the APPswe/PS1dE9 mice. However, this hypothesis cannot be confirmed by the current study. Importantly, these high doses of vitamin D have been demonstrated to be safe in a recent human clinical trial [26].

The current study supports the idea that memantine combined with high dose vitamin D enrichment may provide benefits with respect to gait in people with AD. Given the high accessibility and low cost of vitamin D supplements, the potential benefits of treatments incorporating vitamin D supplementation should be further explored in clinical studies.

### Effect of Age

It is also worthwhile to examine the role of age in the spatiotemporal gait pattern of the APPswe/PS1dE9 mouse model. When comparing control mice to vitamin D-deficient mice from month 6 onwards, age had a significant effect on swing, stride length regularity, and swing speed. Similarly, when comparing the vitamin D deprived groups from month 9 onwards, age had a significant effect on swing, swing speed, and swing speed regularity. Note that a longitudinal analysis of all four treatment groups at once was not performed to avoid confounding treatment (diet scheme) with age. Overall, age tended to increase swing and decrease swing speed, but these tendencies did not become apparent until later in the study, when comparing months 9 and 12 to month 15. The effects of age on stride length regularity and swing speed regularity in the APPswe/PS1dE9 mice were less clear.

Human studies of AD have shown progressive alterations in gait over time that include reduced stride length, increased stride length variability, decreased swing, increased swing variability, decreased swing speed, and increased stride width, among others [3]. While a progressive decrease in swing speed aligns with the current study’s findings, the accompanying decrease in swing and stride length suggests a shuffling gait. In contrast, we found an increase in swing and no change in stride length, suggesting that APPswe/PS1dE9 mice do not (at least within the timeline of this study) exhibit a shuffling gait, but rather a slowing of pace.

With respect to the gait parameters for which an effect of age was not observed in this study, the discrepancy with human studies may be due to the timeline of the current study rather than the ability of APPswe/PS1dE9 mice to model spatiotemporal gait. In previous studies of APPswe/PS1dE9 mice ranging from 3 to 12 months old, performance on wire-hanging, rotarod, and balance beam tests were found to worsen with age [22, 23]. It was therefore expected that spatiotemporal measures of gait in this model would also worsen with age and that any changes would be observable within our study’s timeline (6 to 15 months of age). However, to our knowledge, this study is the first to examine spatiotemporal gait parameters in the APPswe/PS1dE9 model. Future studies following APPswe/PS1dE9 mice for longer periods of time might reveal gait changes in older mice that were not captured by the current study and could verify the tendency of swing and swing speed to slow with age beyond month 15.

Lastly, it is important to mention that even healthy C57BL/6 mice have shown slowing of swing speed, increased stride width, and decreased stride length with age within a similar time frame to the current study (4 to 17 months of age) [37]. Though similar changes in stride width and stride length with age were not seen in our APPswe/PS1dE9 mice, this does not dismiss APPswe/PS1dE9 as a model of gait in AD. We did not compare APPswe/PS1dE9 mice to wild-type control mice in this study, and it is entirely possible that APPswe/PS1dE9 mice would show worsened spatiotemporal gait at all time points relative to wild-type controls.

### Limitations

There are several limitations not yet discussed in this study that should be considered when interpreting the results. Regarding the Catwalk system, there was no adaptation training for the mice prior to the CatWalk tests, such that learning may have affected the results. Furthermore, the mice were not provided with motivation to cross the track, such as a shelter placed at the other end of the track. Therefore, our procedure was more akin to open-field testing in which mice may be less consistent in speed. This approach may have resulted in higher variation of stride length, swing, and swing speed than had the mice been motivated. Additionally, the APPswe/PS1dE9 mouse model does not develop neurofibrillary tangles. Future work should examine the potential effects of vitamin D deficiency and treatment with the combination of memantine and vitamin D on spatiotemporal gait in a mouse model that does exhibit tauopathy, in addition to studies in people with AD. Finally, there are limitations inherent in using transgenic mouse models of AD, such as the APPswe/PS1dE9 model, as these models use specific genetic mutations that are implicated in familial Alzheimer’s disease. Therefore, caution must be used when generalizing findings in these models to sporadic Alzheimer’s disease.

## CONCLUSIONS

In this longitudinal study of APPswe/PS1dE9 mice, vitamin D deprivation led to an increase in hind stride width by month 15, with no changes in swing or swing regularity, stride length or stride length regularity, and swing speed or swing speed regularity. Memantine combined with high dose vitamin D, but not memantine alone, prevented the increase in hind stride width. Therefore, we conclude that vitamin D deprivation may lead to impaired postural control, while the combination of memantine and vitamin D enrichment has the potential to mitigate gait changes associated with Alzheimer’s disease.

## ACKNOWLEDGEMENTS

This study was funded by the Schulich School of Medicine and Dentistry, University of Western Ontario, Canada, the Canadian Institutes of Health Research (MD/PhD Studentship, Foundation Grant FDN 148474), and the Research Centre on Autonomy and Longevity, University Hospital of Angers, France.

