## Supplemental Materials for "Effects of memantine and high dose vitamin D on gait in the APP/PS1 mouse model of Alzheimer’s disease following vitamin D deprivation"

### SUPPLEMENTARY MATERIALS

*Gait Analysis – Longitudinal Effects of Memantine & Vitamin D on Vitamin D-Deficient Mice*

Presented here are the results of longitudinal analyses of the three vitamin D deprived groups. Measurements made at month 9 act as a baseline for these groups, as all three groups had been fed the same vitamin D deficient diet up until this time point. Note that a longitudinal analysis of all four groups at once was not performed to avoid confounding the effect of age with the effect treatment (diet scheme).

Presented first are swing, stride length, swing speed and their coefficients of variation (Suppl Fig 1). The linear mixed model for swing revealed a significant effect of age ($F(1,63) = 9.00$, $p < 0.001$), but not treatment ($F(2,37) = 0.24$, $p = 0.79$) or age-by-treatment interaction ($F(4,67) = 0.70$, $p = 0.60$). Post-hoc comparisons tests showed that swing increased significantly from month 9 to month 15 in both the VitD- ($p < 0.05$) and the Mem & VitD+ ($p < 0.05$) groups. Therefore, memantine and vitamin D enrichment did not modify swing, however, age tended to increase it. The linear mixed model for swing CoV revealed non-significant effects of age ($F(1,65) = 0.56$, $p = 0.57$), treatment ($F(2,37) = 1.19$, $p = 0.31$) and age-by-treatment interaction ($F(4,67) = 2.48$, $p = 0.052$). Therefore, memantine and vitamin D enrichment did not modify swing or swing CoV, however, swing tended to increase with age.

The linear mixed model for stride length revealed no significant effects of age ($F(1,64) = 1.31$, $p = 0.28$), treatment ($F(2,37) = 0.56$, $p = 0.58$), or age-by-treatment interaction ($F(4,67) = 0.85$, $p = 0.50$). Memantine and vitamin D enrichment did not affect stride length, nor did stride length change as the mice aged. The linear mixed model for stride length CoV revealed a significant effect of age ($F(1,65) = 6.36$, $p < 0.01$) and age-by-treatment interaction ($F(4,67) = 3.65$, $p < 0.01$), but not treatment ($F(2,37) = 0.19$, $p = 0.83$). Post-hoc comparison tests showed that stride length CoV significantly decreased from months 12 to 15 in the Mem & VitD- group ($p < 0.01$). At month 12, the stride length CoV of the Mem & VitD+ group was significantly lower than that of both the Mem & VitD- ($p < 0.05$) and VitD- ($p < 0.05$) groups. By month 15, however, there were no significant differences between groups, and overall trends with age were not clear.

The linear mixed model for swing speed revealed a significant effect of age ($F(1,65) = 7.23$, $p < 0.01$), but not treatment ($F(2,37) = 0.69$, $p = 0.51$) or age-by-treatment interaction ($F(4,67) = 0.54$, $p = 0.71$). Swing speed tended to decrease with age, particularly in the VitD- and Mem & VitD+ treatment groups. Post-hoc tests showed that swing speed significantly decreased from month 9 to month 15 in the VitD- group ($p < 0.05$). Memantine and vitamin D enrichment did not affect swing speed. The linear mixed model for swing speed CoV revealed a significant effect of age ($F(1,62) = 3.86$, $p < 0.05$), but not treatment ($F(2,37) = 0.54$, $p = 0.59$) or age-by-treatment interaction ($F(4,67) = 1.83$, $p = 0.13$). Therefore, memantine and vitamin D enrichment did not affect swing speed regularity in vitamin D deficient mice. While post-hoc comparison tests showed that swing speed CoV significantly decreased from month 12 to month 15 in the Mem & VitD- group ($p < 0.05$), trends in swing speed CoV with age were not clear.

Finally, front and hind stride widths were considered (Suppl Fig 2). The linear mixed model for front stride width revealed there were no significant effects of age ($F(1,49) = 3.35$, $p = 0.06$), treatment ($F(2,37) = 0.55$, $p = 0.58$), or age-by-treatment interaction ($F(4,67) = 0.07$, $p = 0.99$). Likewise, the linear mixed model for hind stride width revealed there were no significant effects of age ($F(1,55) = 1.28$, $p = 0.28$), treatment ($F(2,37) = 0.57$, $p = 0.57$), or age-by-treatment interaction ($F(4,67) = 1.33$, $p = 0.27$). These results indicate that neither memantine and vitamin D enrichment nor age modified front or hind stride width in vitamin D deficient mice.

**Supplementary Figure 1. Effect of memantine plus vitamin D on swing, stride length, and swing speed of vitamin D deficient mice:** The CatWalk gait analysis system was used to measure spatiotemporal parameters of gait in APPswe/PS1dE9 mice. All groups were fed a vitamin D-deficient diet starting at six months. Treatment began for the Mem & VitD- and Mem & VitD+ groups at nine months and continued for the duration of the study. Shown here are swing (A), stride length (B), swing speed (C) and their respective coefficients of variation (D-F) in the Mem & VitD-, Mem & VitD+, and VitD- treatment groups at 9, 12, and 15 months of age. Mem, memantine; VitD+, vitamin D-enriched; VitD-, vitamin D-deficient. **p* < 0.05, ***p* < 0.01, [a]VitD- vs. Mem & VitD+: *p* < 0.05, [b]Mem & VitD- vs. Mem & VitD+: *p* < 0.05.

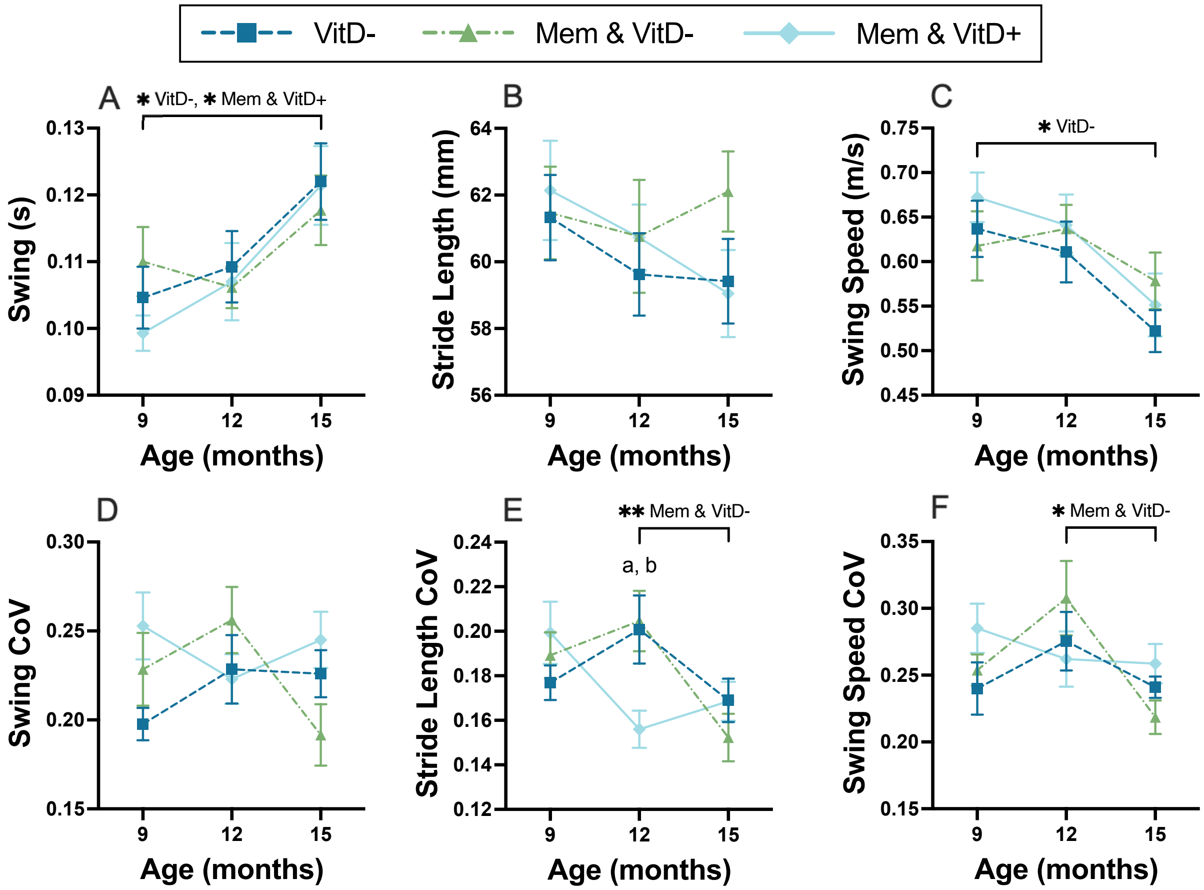

**Supplementary Figure 2. Effect of memantine plus vitamin D on front and hind stride width of vitamin D deficient mice:** The CatWalk gait analysis system was used to measure spatiotemporal parameters of gait in APPswe/PS1dE9 mice. All groups were fed a vitamin D-deficient diet starting at six months. Treatment began for the Mem & VitD- and Mem & VitD+ groups at nine months and continued for the duration of the study. Shown here are front (A) and hind (B) stride width of the Mem & VitD-, Mem & VitD+, and VitD- treatment groups at 9, 12, and 15 months of age. Mem, memantine; VitD+, vitamin D-enriched; VitD-, vitamin D-deficient.

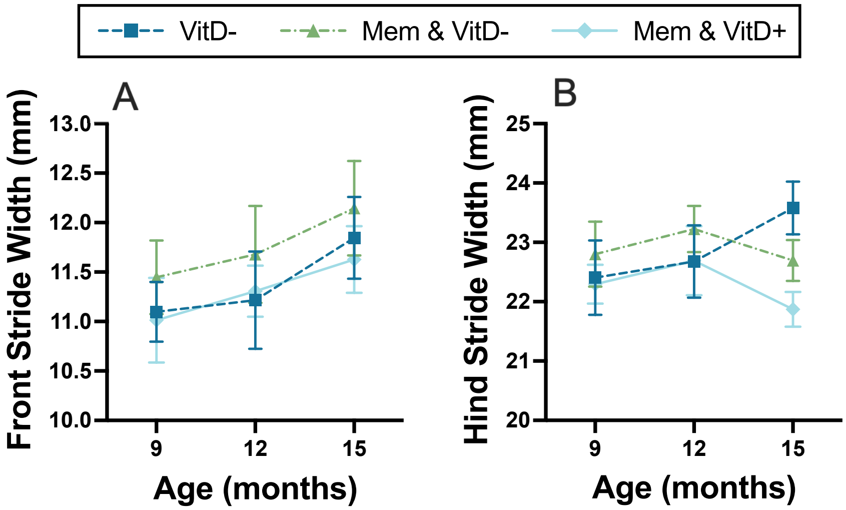
